# Systemic in utero gene editing as a treatment for cystic fibrosis

**DOI:** 10.1101/2024.09.04.611031

**Authors:** Adele S. Ricciardi, Christina Barone, Rachael Putman, Elias Quijano, Anisha Gupta, Richard Nguyen, Hanna Mandl, Alexandra S. Piotrowski-Daspit, Francesc Lopez-Giraldez, Valerie Luks, Mollie R. Freedman-Weiss, James Farrelly, Samantha Ahle, Peter M. Glazer, W. Mark Saltzman, David H. Stitelman, Marie E. Egan

## Abstract

In utero gene editing has the potential to modify disease causing genes in multiple developing tissues before birth, possibly allowing for normal organ development, disease improvement, and conceivably, cure. In cystic fibrosis (CF), a disease that arises from mutations in the cystic fibrosis transmembrane conductance regulator (*CFTR*) gene, there are signs of multiorgan disease affecting the function of the respiratory, gastrointestinal, and reproductive systems already present at birth. Thus, treating CF patients early is crucial for preventing or delaying irreversible organ damage. Here we demonstrate proof-of-concept of multiorgan mutation correction in CF using peptide nucleic acids (PNAs) encapsulated in polymeric nanoparticles and delivered systemically in utero. In utero editing was associated with sustained postnatal CFTR activity, at a level similar to that of wild-type mice, in both respiratory and gastrointestinal tissue, without detection of off-target mutations in partially homologous loci. This work suggests that systemic in utero gene editing represents a viable strategy for treating monogenic diseases before birth that impact multiple tissue types.

## Introduction

In cystic fibrosis (CF), the onset of multiorgan pathology begins during fetal development. CF was first described in infants as a condition that resulted in failure to thrive from lethal malabsorption caused by an abnormal pancreas^1^. We now know that upwards of 85% of children display exocrine pancreatic insufficiency before the age of one^2–4^. In addition to pancreatic dysfunction, children with CF exhibit signs of multiorgan disease at the time of birth including reduced birth weight^5^, meconium ileus and subsequent microcolon (15-20%)^6^, tracheomalacia (6-41%)^7,8^, biliary cirrhosis (11%)^9^, and 95% of males have congenital bilateral absence of the vas deferens, which leads to issues with fertility. There is also significant respiratory pathology that ultimately drives morbidity and mortality in patients with CF, but these pulmonary manifestations are quite limited in the neonatal period. Given the extent of disease present at birth, and the limited lung pathology in the perinatal period, there is a growing consensus that treating CF patients early is crucial in preventing or delaying irreversible organ damage^10^.

When CF was first described, over 80% of patients died within the first year of birth. With the development of treatments and the establishment of centers with aggressive, comprehensive care, the median survival of patients with CF continues to increase. For many patients, the current treatment paradigm includes CFTR modulators, which function by rescuing endogenous CFTR protein activity and have shifted the focus of CF treatment away from supportive care. Although a breakthrough in treatment, modulators are taken daily, require lifelong treatment, are expensive, and are not available for all *CFTR* mutations. While the currently available treatment greatly improves the quality of life and life expectancy for many patients with CF, none of these therapies correct the underlying genetic defect, that if achieved in the correct organs and cell types, could result in cure.

Systemic (i.e. intravenous) delivery of gene editing therapeutics holds great promise in the treatment of diseases that affect multiple organs, such as CF. Recent studies have demonstrated the potential to correct CF disease-causing mutations via the intravenous (IV) delivery of nanoparticles (NPs) containing gene editing technologies in adult animals^11–13^. These studies include our recent work where we first demonstrated the potential for multi-organ editing in CF after systemic delivery of polymeric nanoparticles loaded with peptide nucleic acids (PNAs) and single-stranded donor DNA – a non-nuclease mediated gene editing approach that has been shown to have extremely low to undetectable off-target effects due to the lack of inherent nuclease activity^14^ – in postnatal animals^11^. Treatment prior to birth, during or before the onset of pathology, has yet to be demonstrated.

Given the extent of disease present at birth, we hypothesized that *CFTR* mutation correction in utero could prevent irreversible organ damage, increase functional CFTR activity, or perhaps cure an individual of CF. Previously, we have used a similar approach for gene correction in β-thalassemia in mice, where a single in utero IV injection of PNA/DNA NP treatment led to phenotypic and genomic changes that persisted into adult life^15^. Here, we demonstrate that NPs loaded with PNA and donor DNA can successfully edit the developing fetal lung in a GFP reporter mouse model and that PNA/DNA NP treatment can correct the F508del mutation in utero, resulting in sustained postnatal CFTR function in tissue types affected by CF after a single in utero NP treatment.

## Results

### Nanoparticle uptake and tropism in the fetal lung

We previously established the capacity to deliver polymeric NPs formulated from poly(lactic-co-glycolic acid) (PLGA) to numerous fetal tissues, including the lung and gastrointestinal tract, using two different in utero delivery approaches: IV administration via the vitelline vein or intra-amniotic (IA) administration via direct injection into the amniotic fluid^15^. Because the respiratory tract is the main source of pathology in patients affected by CF, we first sought to determine if either of these approaches led to more robust intracellular NP uptake in the lung and to explore the cellular tropism of both delivery techniques.

We administered NPs containing the green fluorescent dye, DiO, to fetal mice either IV on embryonic day 15.5 (E15.5) or IA on E16.5 – the earliest gestational age in which we have seen accumulation of NPs in the fetal lung (Fig. 1a)^15^. We compared the fetal lung tropism of PLGA and a formulation containing a blend of PLGA and poly(beta amino ester) (PBAE) surface modified with the cell-penetrating peptide MPG, which has previously been shown to result in higher uptake and CFTR editing in airway epithelial cells^14,16^. When characterized by dynamic light scattering (DLS) and scanning electron microscopy (SEM), both NP formulations were spherical and approximately 250 nm in diameter; they differed in zeta potential, with PLGA NPs carrying a negative charge, compared to the positively charged PLGA/PBAE/MPG particles (Supplementary Table 1 & Supplementary Fig. 1).

**Figure 1.**
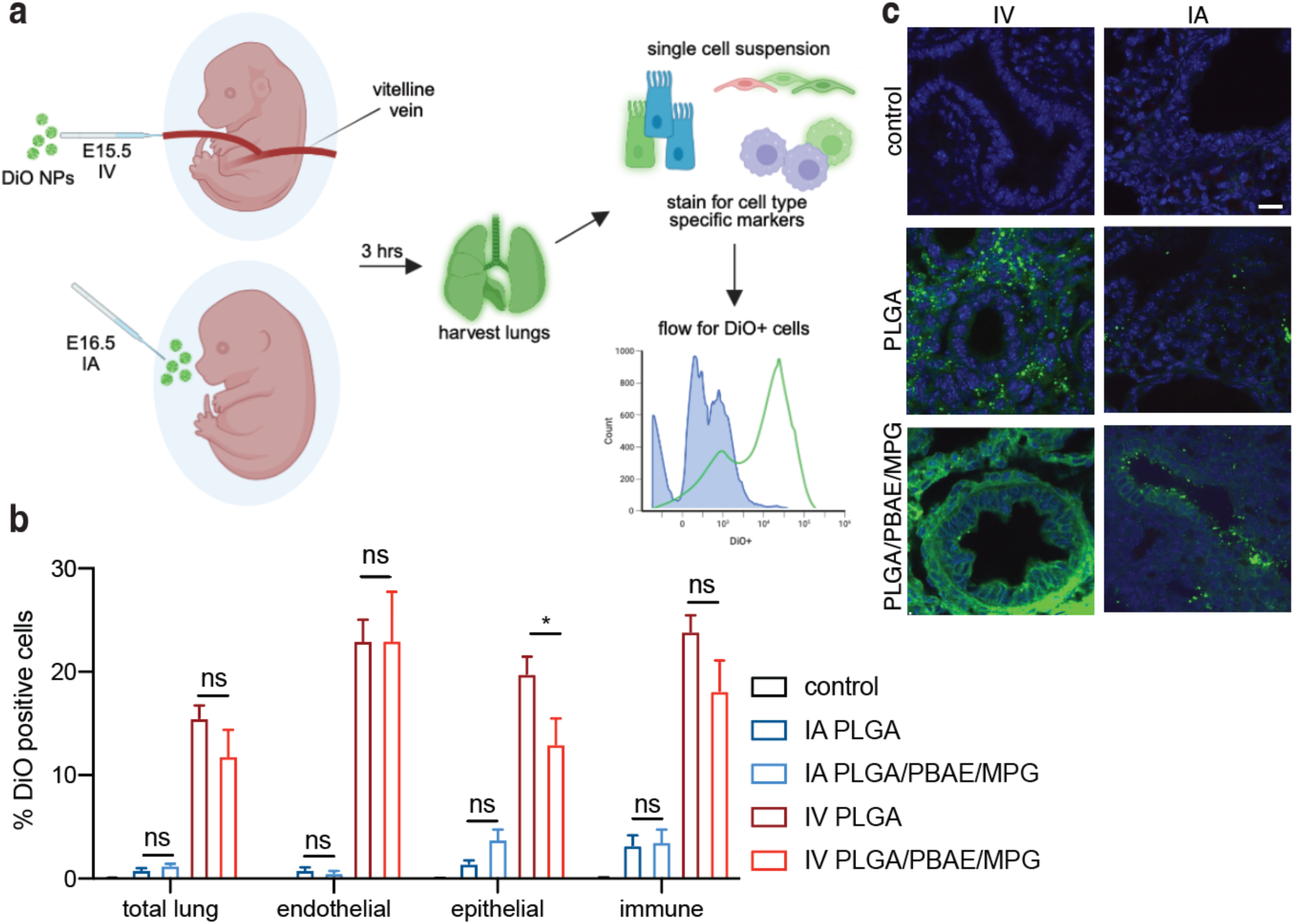
In utero delivery of nanoparticles to the lung. a, schematic of E15.5 intravenous (IV) and E16.5 intra-amniotic (IA) nanoparticle (NP) delivery routes via micropipette injection into the vitelline vein or directly into the amniotic fluid and analysis. b, lung cellular tropism of DiO loaded PLGA and DiO loaded PLGA/PBAE/MPG NPs (% DiO positive cells) 3h after either E16.5 IA or E15.5 IV delivery, n=5 for each treatment group, data are mean ± s.e.m., statistical analysis by two-way ANOVA, *P<0.03. c, confocal images of the distribution of DiO loaded NPs (green) 3h after either E16.5 IA (top) or E15.5 IV (bottom) delivery with images of uninjected age matched controls to the left. Nuclei within the tissues were stained with Hoescht (blue), scale bar = 10 μm.

When delivered in utero, both formulations resulted in NP accumulation in the fetal lung three hours after either IV or IA delivery, with IV delivery resulting in a significantly higher percentage of cells containing NPs when evaluated via flow cytometry (Fig. 1b) or confocal microscopy (Fig. 1c). Using flow cytometry we also assessed the tropism of the NPs for specific populations of cells in the lung including endothelial (CD31+), epithelial (EpCAM+), and immune (CD45+) cells. We found that in utero IV delivery of NPs resulted in particle accumulation in all three cell types, including epithelial cells (Fig. 1b). Other than IV NP delivery to epithelial cells, where PLGA NPs result in significantly higher NP delivery than the PLGA/PBAE/MPG formulation, we found no significant difference between the tropism of the two particle types after IV or IA delivery (Fig. 1b).

IA administration resulted in lower NP accumulation than we expected; to examine the stability of NPs in this setting, we incubated both NP formulations in murine amniotic fluid collected from E16.5 fetuses and measured the change in size and polydispersity index (PDI) using successive DLS measurements over an hour. Both formulations were unstable in murine amniotic fluid when compared to incubation in PBS, as evidenced by increasing and vacillating size and PDI during this time frame (Supplementary Fig. 2).

### Gene editing of fetal lungs

We next sought to determine if NP accumulation within the lung after IV or IA NP delivery correlated with the ability to edit fetal lung cells (Fig. 2a). To assess gene editing efficiency, we used a transgenic reporter mouse that contains an eGFP gene that has been disrupted by an intron containing a splice-site mutation that results in aberrant splicing, retention of a segment of the intron, and abrogation of eGFP fluorescence^17^. Correction of the splice-site mutation results in correct intronic splicing, allowing for eGFP fluorescence that can be used as a readout for gene editing (Fig. 2b). To achieve gene editing we used NPs loaded with triplex-forming PNAs and single-stranded donor DNAs.

**Figure 2.**
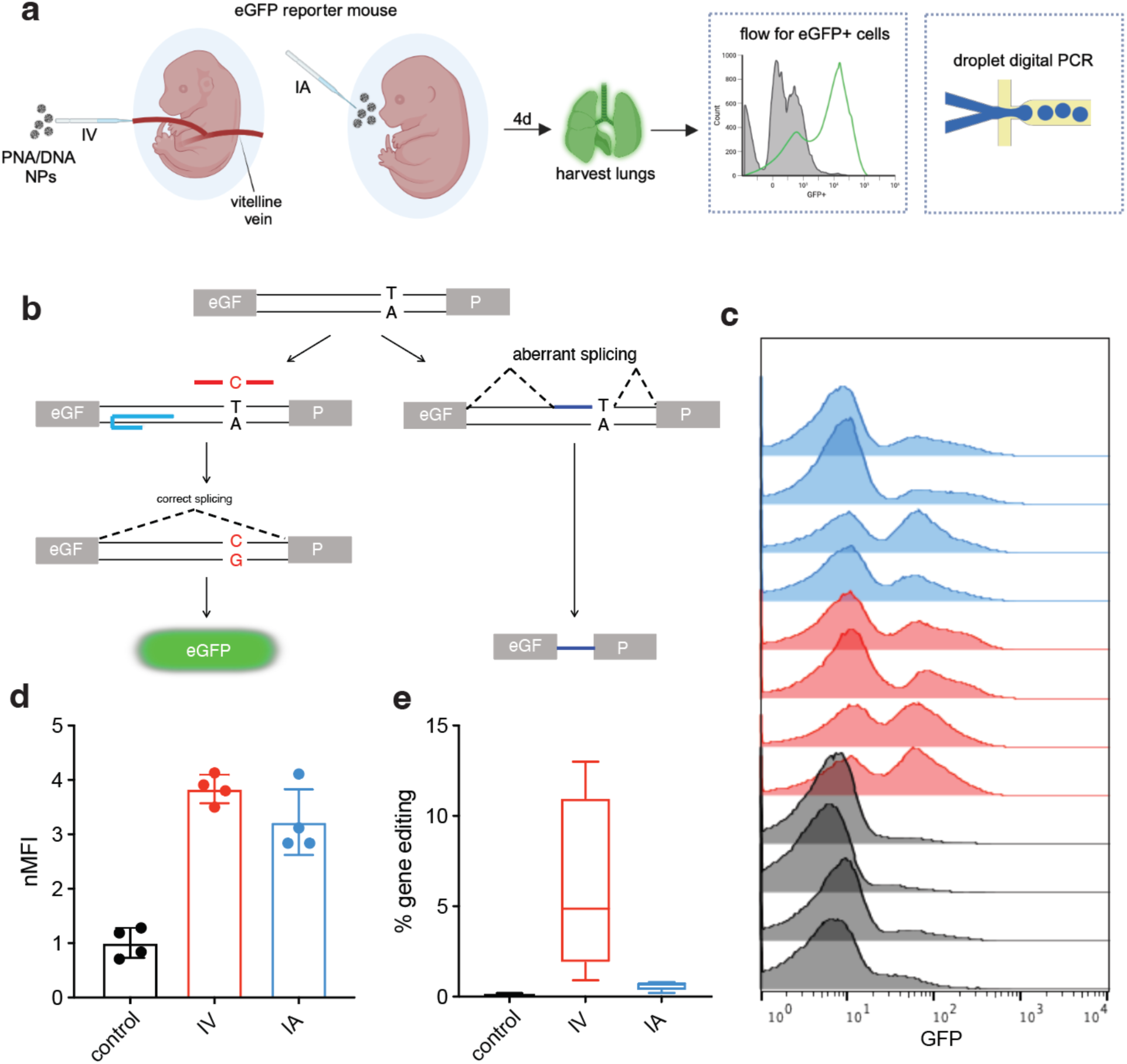
Gene editing of fetal lungs in an eGFP reporter mouse model after in utero nanoparticle delivery. a, schematic of in utero NP delivery and analysis. b, schematic of the eGFP reporter mouse model that contains a mutation in an intron that interrupts the eGFP coding sequence (grey boxes). The mutation creates an aberrant splice site that leads to retention of an intronic fragment in the spliced mRNA, preventing proper translation of eGFP (right). Correction of the splice-site mutation with PNA (blue) and donor DNA (red) designed to target the mutation, results in correct splicing and restores eGFP expression (left). c-d, flow cytometry histograms (c) and normalized mean fluorescence intensity (nMFI) of the total lung cell population four days after either intravenous (IV, red) or intra-amniotic (IA, blue) nanoparticle (NP) treatment compared to untreated lungs (grey), n=4 for each treatment group, data are mean ± s.e.m. e, the frequency of gene editing was measured in genomic DNA (gDNA) derived from fetal lungs four days after in utero NP treatment by droplet digital PCR (ddPCR), n=4 for each treatment group, the horizontal lines within the boxes indicate the median, the box indicates the first and third quartile, and the whiskers represent the range.

To assess in utero gene editing in this reporter mouse we used a PNA designed to target a homopurine stretch upstream of the splice-site mutation^15,18^. This PNA contained a tail clamp (tc) designed to extend the Watson-Crick binding domain to increase binding affinity^19,20^ and a mini-polyethylene glycol side chain at the γ-backbone position, which has been shown to increase solubility and binding affinity for the DNA target^21^. This γtcPNA was encapsulated in PLGA NPs with a 60-bp donor DNA that contains the correct base sequence in a 2:1 molar ratio. The NPs were spherical with sizes and zeta potentials consistent with previous formulations (Supplementary Fig. 1 & Supplementary Table 1). PNA/DNA NPs were delivered IV on E15.5 or IA on E16.5. Four days after treatment, fetal lungs were harvested and dissociated into single cells. eGFP fluorescence was assayed by flow cytometry, which revealed a significant increase in eGFP expression after both IA and IV in utero NP treatment (Fig 2c&d). The frequency of gene correction was measured by droplet digital PCR (ddPCR). As predicted by the cellular tropism measured with fluorescent NPs, IV NP treatment resulted in a significantly higher level of gene editing (∼6%) in the lung than that achieved by IA NP treatment (∼0.6%) (Fig. 2e).

### Functional CFTR activity after in utero gene editing in a mouse model of cystic fibrosis

Next, we investigated if PNA/DNA NPs could be used to correct the most common disease causing CFTR mutation, F508del, after in utero delivery in a mouse model homozygous for the 3 base pair deletion (Fig. 3a). PNA and donor DNA designed to correct the F508del mutation in mice^14^ were loaded into PLGA NPs. Again, the NPs displayed a spherical morphology, with size and zeta potential consistent with previous formulations (Supplementary Fig. 1 & Supplementary Table 1)^14^. A single dose of PNA/DNA NPs were administered to F508del CF mouse fetuses IV on E15.5 and IA on E16.5. Eight months after fetal treatment a nasal potential difference (NPD) assay was performed to measure the activity of CFTR in the nasal epithelium of treated (both IA and IV) and untreated mice. The NPD is a non-invasive assay used to detect chloride transport in vivo. In this assay, the lack of activation of cyclic AMP-stimulated chloride efflux with forskolin (FSK) is the hallmark of CF affected epithelia and the restoration of this response serves as a surrogate of CFTR activity. After in utero PNA/DNA NP treatment we found that the impaired response to cyclic AMP stimulation was corrected after both IA and IV NP treatment and more robustly corrected, to a value similar to that of wild-type mice, after IV NP treatment (Fig. 3b). Representative NPD traces are displayed in Supplementary Fig. 3. We additionally measured the rectal potential difference (RPD) for the mice who received IV NP treatment and found a significant improvement in the RPD, to a level similar to that of wild-type mice (Fig. 3c).

**Figure 3.**
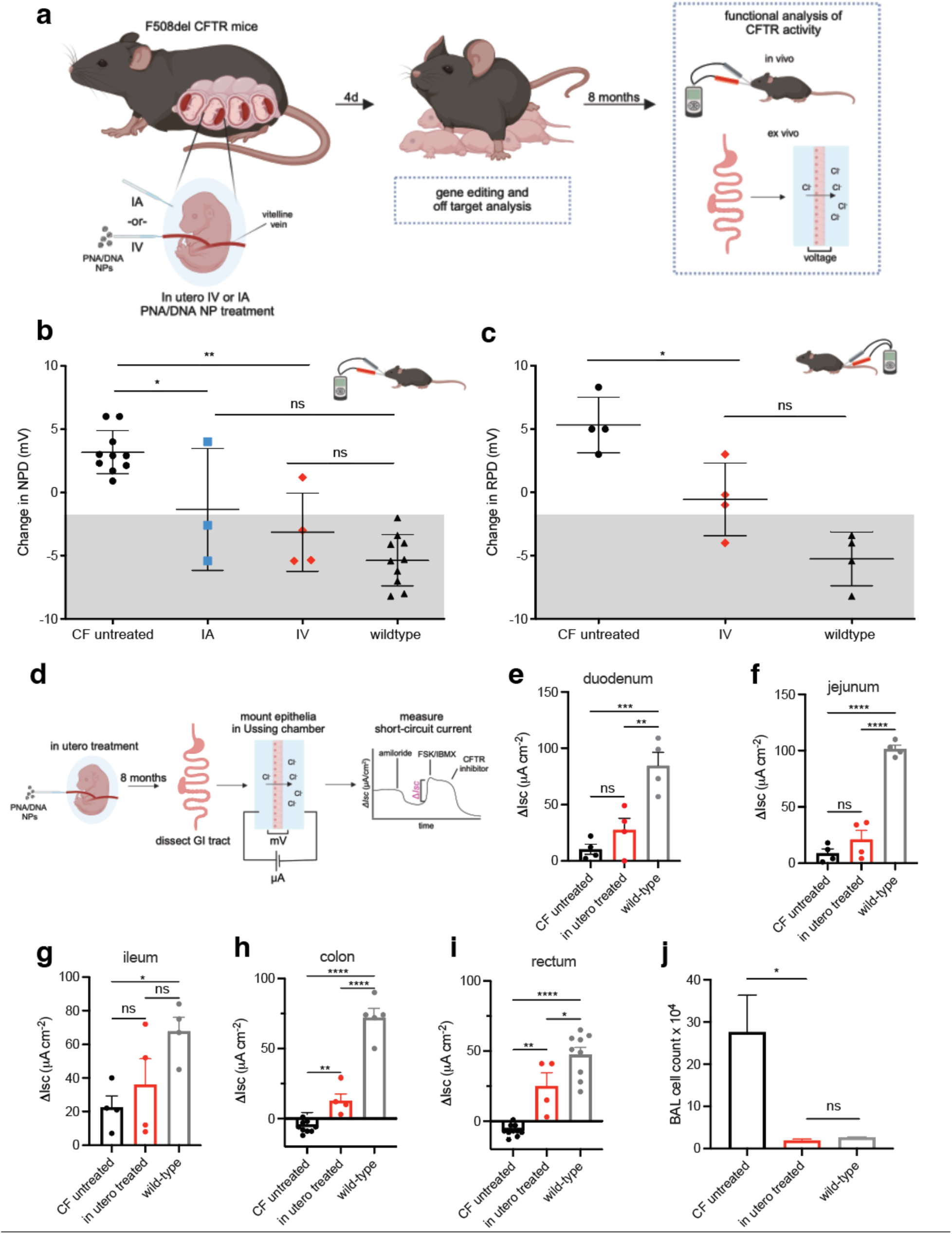
Phenotypic disease improvement in cystic fibrosis (CF) adult mice after in utero gene editing. a, schematic of in utero nanoparticle (NP) treatment and analysis. b, the nasal potential difference (NPD) and c, rectal potential difference (RPD) in untreated CF mice (NPD n=12, RPD n=4), wild-type mice (NPD N=7, RPD n=4) and CF mice eight months after intra-amniotic (IA)(NPD n=3), or intravenous (IV)(NPD n=4, RPD n=4) in utero nanoparticle treatment. Each mouse is represented by an individual data point, the mean is shown with a horizontal line, surrounded by error bars showing the s.e.m. d-i, schematic of ex vivo short circuit (βI_sc_) analysis using the Ussing chamber assay (d) for the duodenum (e), jejunum (f), ileum (g), colon (h), and rectum (i). j, bronchoalveolar lavage (BAL) fluid cell counts eight months after IV in utero NP treatment (red) compared to untreated CF (black) and wild-type (grey) mouse controls, data are mean +/- s.e.m. Statistical analysis by one way ANOVA, *P<0.05, **P<0.01, ***P<0.001, ****P<0.0001.

In addition to the in vivo phenotypic correction, we assessed CFTR function ex vivo in epithelial gastrointestinal tissue eight months after systemic fetal PNA/DNA NP treatment using an Ussing chamber assay that measures CFTR mediated ion transport as a change in short-circuit current (Δ*I_sc_*) in response to a CFTR-stimulating cocktail of FSK and 3-isobutyl-1-methylxanthine (IBMX) (Fig. 3d). We found a significant increase in the Δ*I_sc_* in the colon and rectum in IV treated mice when compared to untreated CF mice (Fig 3h&i). We observed a more modest increase in the Δ*I_sc_* in the more proximal GI tissue including the duodenum, jejunum, and ileum (Fig. 3e-g).

We further demonstrated phenotypic improvement in our in utero treated mice by measuring cell counts in the bronchoalveolar lavage (BAL) fluid of mice, which typically contains immune cells such as alveolar macrophages and neutrophils. When compared to untreated CF mice, we observed significantly lower cell counts in our in utero treated mice eight months after birth. There was no significant difference between the cell counts in our treated mice when compared to wild-type mice, which suggests decreased inflammation in the airway secondary to restoration of CFTR function (Fig. 4j). Taken together, we demonstrate long-term functional disease improvement in both respiratory and gastrointestinal tissues after a single in utero PNA/DNA NP treatment.

**Figure 4.**
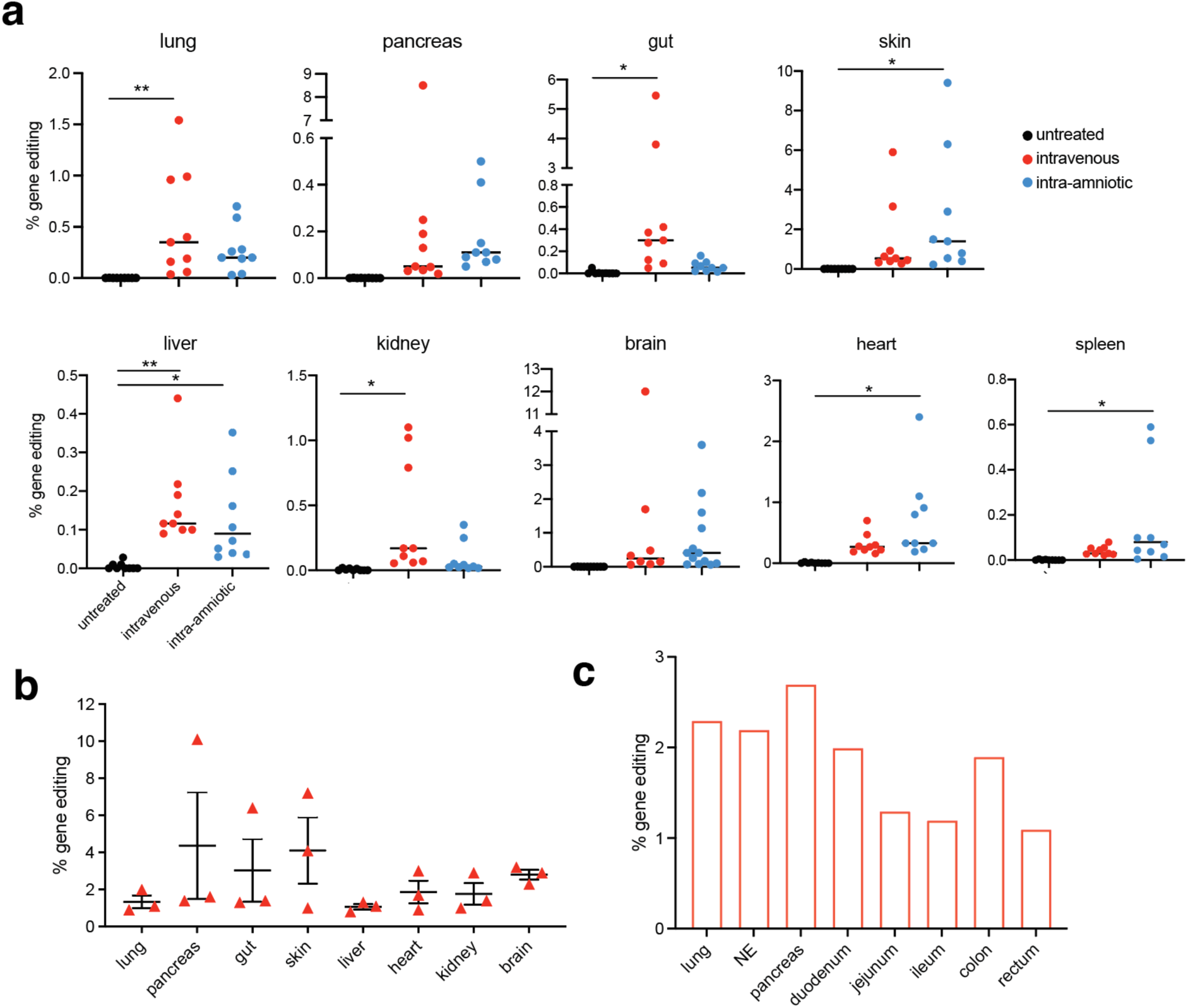
Gene editing frequency in multiple organs after in utero gene editing in cystic fibrosis (CF) mice. a, droplet digital PCR (ddPCR) quantification of gene editing in genomic DNA (gDNA) from organ biopsies four days post in utero intravenous (IV) or IA (intra-amniotic) PNA nanoparticle (NP) treatment compared to untreated CF controls, (n=9, data are mean ± s.e.m). b, deep sequencing quantification of gene editing in gDNA from organ biopsies four days post IV PNA NP treatment, n=3, data are mean ± s.e.m. c, deep sequencing quantification of gene editing in an adult mouse eight months after IV NP treatment, NE = nasal epithelium. Statistical analysis by one way ANOVA, *P<0.05, **P<0.01, ***P<0.001, ****P<0.0001.

Due to limitations in the potential difference assays, the experiments cannot be reliably performed until the mice are 20g. Fortuitously, this allowed us to examine the longevity of gene editing achieved after a single in utero NP treatment dose. We observed improved CFTR function to at least eight months of age, which contrasts with the NPD response seen in adult mice treated systemically with PNA/DNA NPs, where the change in NPD is attenuated after four weeks and repeat dosing is required to sustain functional CFTR activity^11^.

We performed our PNA-NP treatments during the transition between the pseudoglandular and canalicular phases of lung development, in which the lung is rapidly dividing to generate the lung tissue necessary to sustain postnatal life^22^. This environment may be particularly primed for DNA repair, and we hypothesized that the developing fetal lung may be a more hospitable environment for PNA-mediated gene editing than the adult lung. Using RNAseq, we evaluated the expression of genes involved in pathways related to gene editing: the HDR, NER, MMR and FA DNA repair pathways^23–27^. We found that genes in these pathways are elevated in E16.5 fetal compared to adult lung tissue (Supplementary Fig. 4). When we analyzed a larger list of genes associated with DNA repair, 162 of the 226 genes involved with DNA repair show significant differences in expression in the fetal and adult lungs, with the majority of DNA repair genes having higher expression in the fetal lung (Supplementary Fig. 5).

### Genotypic correction after PNA-mediated in utero gene editing

We next sought to quantify the level of gene editing that led to the observed phenotypic CFTR correction. The treated fetal mice were born an average of four days after treatment and a droplet digital PCR (ddPCR) assay designed to discern between the edited *CFTR* allele, containing a CTT insertion, and the mutant F508del allele was used to measure the percent of gene editing in numerous fetal tissues at this time (Fig. 4a). We observed gene correction in numerous tissues relevant to CF after systemic NP treatment including: lung (up to 1.5%), pancreas (up to 8.5%), gut (up to 5.5%), skin (up to 6%), and liver (up to 0.4%). While less relevant to CF, we also found editing in the kidney (up to 1.2%) and brain (up to 12%). A lesser extent of editing was also found in many tissues after IA NP treatment including: lung (up to 0.7%), gut (up to 0.2%), pancreas (up to 0.5%), brain (up to 3.6%), liver (up to 0.4%), skin (up to 9.4%), heart (up to 1.1%), and spleen (up to 0.6%) (Fig. 4a). The level of gene correction after fetal IV NP treatment measured with ddPCR was also confirmed by deep sequencing (Fig. 4b). Deep sequencing was also performed on genomic DNA isolated from a mouse eight months after IV in utero NP treatment, which revealed a sustained level of gene correction in tissues relevant to CF into adulthood (Fig. 4c). Lastly, we used deep sequencing to assess for potential off-target effects in the lung both four days and eight months after PNA/DNA NP treatment by evaluating ten genomic sites with partial homology to the binding site of the PNA and one site with partial homology to the donor DNA^11^. The results revealed an undetectable mutation frequency at these off-target sites in both early postnatal and adult mice (Supplementary Table 2).

## Discussion

CF is a multiorgan disease, with significant manifestations prior to and at the time of birth. Given this extent of disease, it is possible that a prenatal approach to disease treatment may be required to truly cure a patient of CF. We show, for the first time, that cystic fibrosis can be treated before birth by correcting the most common disease-causing mutation, F508del, which results in sustained editing and phenotypic disease improvement in multiple organs affected by CF, including the respiratory and gastrointestinal tracts. Importantly, these effects last into adult life. This result is accomplished by using a single treatment with non-nuclease, triplex forming, PNAs co-delivered with a short donor DNA in polymeric PLGA NPs by either IV injection via the vitelline vein, or by instillation directly into the amniotic fluid, which results in fetal inhalation and ingestion of the NP containing fluid.

PNA/DNA NPs have previously been used to show both phenotypic and genotypic correction of *CFTR* mutations in vitro, in a physiologically relevant model of nasal epithelial cells (NECs) grown at air liquid interface (ALI), and in vivo in adult mice after both topical intranasal and IV delivery^11,14^. A recent study established the feasibility of an IV (i.e. systemic) approach for CF treatment with PNA NPs and also found multiorgan genotypic and phenotypic correction; however, repeat dosing was required to sustain measurable CFTR activity in functional assays^11^. The majority of CF diagnoses result from newborn screening; however, with the advent of non-invasive prenatal genetic testing, such as the sequencing of cell free fetal DNA, CF can now be diagnosed early during gestation. The Cystic Fibrosis Foundation Patient Registry Annual Data Report indicates that the number of prenatal CF diagnoses are increasing each year, with approximately 10% of new diagnoses made before birth in 2021. Early diagnosis opens the window for early intervention. Our work here indicates that there are potential advantages of in utero treatment of CF, as we are able to achieve sustained functional CFTR activity into adulthood of the mouse after a single dose of PNA/DNA NPs in utero using a fraction of the dose needed for similar results in adult mice (approximately 3.2 nmol of PNA were used to treat each adult mouse, and 0.08 nmol of PNA per fetal mouse). The fetal environment is also rife with rapidly dividing stem cell populations, which are the presumed targets that would result in sustained gene correction. We have previously demonstrated that fetal, compared to postnatal, PNA/DNA NP treatment results in a higher population of edited hematopoietic stem and progenitor cells in a mouse model of β-thalassemia^18,28^. As we show here, DNA repair pathways including HDR, NER, MMR, and FA, which are implicated in PNA and CRISPR/Cas9 mediated gene editing, are highly expressed in the murine developing fetal lung compared to the adult lung – a finding that could contribute to the higher frequency of gene correction and longevity of phenotypic response seen in the in utero treated CF mice.

This work examines administration of NPs to the developing murine fetus using two different techniques, both IV and IA. We have previously shown that the administration of PLGA NPs via both in utero techniques in mice is safe, without an impact on survival, growth, or cytokine expression^15^. Here, we show that IV NP administration results in a higher accumulation of NPs in the fetal lung than IA, which correlates to higher levels of editing in the lung in a reporter mouse and improved phenotypic improvement and higher levels of gene editing in the F508del CF mouse. Our work suggests that the PLGA based NPs used for this study may be unstable in murine amniotic fluid, possibly from aggregation or protein deposition, which could diminish their ability to enter cells and deliver nucleic acids. The behavior of NPs in protein-rich biologic fluids is an area of interest for many labs; understanding this behavior is important for the successful translation of many NP-based technologies from lab into clinical practice. It is known that upon contact with biologic fluids, NPs are immediately coated with proteins and other macromolecules^29–32^. This rapid formation of a protein corona influences the subsequent behavior of NPs, including the hydrodynamic size, surface, charge and aggregation^33–35^. Further, the absorbed proteins control NP interaction with cell membranes and potentially alter the mechanism of uptake^36–39^. The corona ends up defining the biological identify of the NPs, and can influence parameters such as biodistribution, toxicity, and internalization^40–42^. The ability to modulate the composition of the biomolecule corona is an exciting area of ongoing investigation and further understanding could improve in vivo delivery in numerous applications. For example, recent work on the inclusion of selective organ targeting (SORT) lipids in lipid nanoparticle (LNP) formulations, which alters the composition of the LNP surface, suggests that chemical changes in LNP composition affect the tissue tropism for systemically delivered LNPs, secondary to differences in protein corona formation^43^. Further understanding and optimization of how polymeric NPs perform in amniotic fluid could prove useful as a means to deliver gene editing cargo to pulmonary epithelial cells.

Prior to this work, therapeutic in utero gene editing of a monogenic disease without the use of a viral vector has been demonstrated in only one prior study. In utero gene editing was first shown by our group using PNA/DNA NPs to correct a disease causing β-globin mutation in hematopoietic stem cells in a mouse model of β-thalassemia, which resulted in sustained postnatal elevation of the blood hemoglobin concentration and improved survival with no detected off-target mutations in partially homologous loci. Here, we also show similar long-term phenotypic improvement with undetectable off-target mutations in partially homologous loci. We are limited in terms of our off-target analysis to the investigation of sites with partial homology to the binding sites of our PNA and donor DNA, as this gene editing technique is non-nuclease mediated, making use of endogenous, high fidelity repair pathways. We have shown in multiple ex vivo and in vivo systems that this system results in extremely low to undetectable levels of off-target mutagenesis at sites of partial homology, a property of non-nuclease based gene editing that is particularly attractive when considering modifying the developing fetal genome.

Nuclease-based gene correction technologies such as CRISPR-Cas systems with homology directed repair (HDR) templates, base editors, and prime editors additionally show great promise in the treatment of CF. Two notable recent studies have demonstrated systemic correction of currently untreatable nonsense mutations via delivery of optimized lung SORT LNPs containing cargo for either CRISPR-Cas9 mediated HDR^13^, or base editing^12^. In the study that employed adenine base editing (ABE), correction of the target nucleotide of the R553X mutation was achieved in 50% of lung basal cells ten days after a single ABE-LNP treatment. As the mouse model used does not exhibit pathologic features of CF, organoids were created from intestinal stem cells from the model. After treatment of the organoids with LNPs, 82% of the organoids, with gene correction at a similar level to the basal cells in the mouse model, exhibited swelling, indicative of restored CFTR function^12^. Although the levels of editing observed in our work are lower, we show here that lower levels of gene correction are still capable of producing long-lasting phenotypic disease improvement. We additionally see variable levels of editing in the CF mouse tissue, and going forward, it would be of interest to define the specific cell types edited with each delivery route and if there is any anatomic relationship or variance throughout tissues of interest (i.e. proximal vs distal airways, central vs peripheral liver). Future research into the optimization of polymeric NP formulations and PNA chemistry are also areas of ongoing interest and may prove fruitful in increasing in vivo editing levels^44,45^.

This work provides a foundation for systemic in vivo gene editing at a time during development that results in sustained phenotypic disease improvement after a single NP treatment and could provide a means of preventing permanent organ damage. Of interest, recent clinical case reports have remarkably shown that in utero exposure to modulator therapy may prevent organ dysfunction at the time of birth. For example, there is report of in utero exposure to modulator therapy in two infants homozygous for F508del mutations (one male, one female) that resulted in normal vas deferens in the male and normal pancreatic function in both infants^46^. Additional reports have documented normal pancreas function, a false newborn screen, and successful treatment of meconium ileus^47,48^. These promising findings serve as motivation into future work on in utero gene editing as a treatment for CF, as prenatal gene correction may provide the potential for a one-time cure.

## Methods

### Nanoparticle fabrication and characterization

PNA/DNA NPs were formulated using a double-emulsion solvent evaporation technique modified to encapsulate PNA and DNA oligomers^18,49^. PNAs and donor DNAs were dissolved in 60.8 μl DNAse-free water. All nanoparticle batches had 2 nmole mg^-1^ of PNA and 1nmole mg^-1^ of donor DNA. For PLGA NPs, the encapsulant was added dropwise to a polymer solution containing 80 mg 50:50 ester-terminated PLGA (0.95-1.2 g dl^-1^, LACTEL absorbable polymers; Birmingham, AL) dissolved in dichloromethane (800 μl), then ultrasonicated (3 × 10 s) to formulate the first emulsion. To form the second emulsion, the first emulsion was added slowly dropwise to 1.6 ml of 5% aqueous polyvinyl alcohol and then ultrasonicated (3 × 10 s). This mixture was finally poured into 20 ml of 0.3% aqueous polyvinyl alcohol and stirred for 3 hours at room temperature. Nanoparticles were then thoroughly washed with 20 ml water (3x) and further collected each time by centrifugation (25,644 x *g* for 10 min at 4 °C). Nanoparticles were resuspended in water, frozen at -80 °C, and then lyophilized. Nanoparticles were stored at -20 °C after lyophilization^18^.

PBAE was synthesized by a Michael addition reaction of 1,4-butanediol diacrylate (Alfa Aesar Organics, Ward Hill, MA) and 4,40-trimethylenedipiperidine (Sigma, Milwaukee, WI) as previously reported^50^. DSPE-PEG(2000)-maleimide was purchased from Avanti Polar Lipids (Alabaster, AL). MPG peptides were purchased from Keck (Yale University). The cell penetrating peptide (CPP), MPG, was covalently linked to DSPE-PEG-maleimide as previously reported^16^. PLGA/PBAE particles contained 15% PBAE (wt%), and were prepared as above except for the following two modifications. Solvent from these particles was evaporated overnight in 0.3% polyvinyl alcohol instead of for 3 hours as above. To make surface-modified particles, DSPE-PEG-MPG was added to the 5.0% PVA solution during formation of the second emulsion at a 5 nmol mg^-1^ ligand-to-polymer ratio.

Fluorescent DiO NPs were synthesized using a single-emulsion solvent evaporation technique^46^. DiO was added to the polymer solution at a 0.2% wt:wt dye:polymer ratio. DiO and 80 mg of PLGA (or PLGA/PBAE with 5 nmol mg^-1^ DSPE-PEG-MPG) were dissolved in 800 μl dichloromethane (DCM) overnight. This mixture was added dropwise to 1.6 mL 5% aqueous polyvinyl alcohol (PVA), then ultrasonicated (3 × 10 s) to formulate a single emulsion. This mixture was poured into 20 ml of 0.3% aqueous PVA and stirred for 3 hours (or overnight for PLGA/PBAE NPs) at room temperature. Nanoparticles were then thoroughly washed with 20 ml water (3x) and further collected each time by centrifugation (25,644 x *g* for 10 min at 4 °C). Nanoparticles were resuspended in water, frozen at -80 °C, and then lyophilized. Nanoparticles were stored at -20 °C after lyophilization^18^.

NP morphology was assessed by scanning electron microscopy (SEM) using an XL-30 scanning electron microscope (FEI; Hillsboro, Oregon) as previously described^51^. Prior to imaging with SEM, samples were coated with 25 nm-thick gold using a sputter coater. Dynamic light scattering (DLS) was performed to measure the NPs hydrodynamic diameter and zeta potential using a Malvern Nano-ZS (Malvern Instruments, UK).

### Mouse models

All animal use was in accordance with the guidelines of the Animal Care and Use Committee of Yale University and conformed to the recommendations in the Guide for the Care and Use of Laboratory Animals (Institute of Laboratory Animal Resources, National Research Council, National Academy of Sciences, 1996). C57BL/6 mice were obtained from Charles River Laboratories (Wilmington, MA). The eGFP transgenic mouse model was obtained from Dr. Ryszard Kole, University of North Carolina, Chapel Hill^17^.

The CF mouse model contains a F508del mutation on a fully backcrossed C57/BL6 background^52^. The genotype of all animals used for this study is indicated in the text. The CF mice were genotyped as follows. Tail or ear tissue was collected from pups between P10-P14 and digested overnight at 55 °C in 600 μl of TNE buffer (100 mM EDTA, 400 mM NaCl, 0.6% SDS, and 10 mM Tris) and 20 μl of proteinase K (10 mg ml^-1^). Next, 166 μl of 6M NaCl was added to the digested tissue, the solution was mixed and then centrifuged at maximum speed at room temperature for 20 minutes. The supernatant was transferred to a new tube and 800 μl of 100% ethanol was added and the sample was mixed by inversion, followed by centrifugation at maximum speed for 10 minutes at room temperature. The supernatant was discarded and the DNA pellet is washed with 70% ethanol. The pellet was re-centrifuged at maximum speed for five minutes at room temperature and the supernatant is again discarded. The pellet was allowed to dry fully before being resuspended in 250 μl dH_2_O.

Genotyping PCR was performed using the following primers: forward – 5’-GAGTGTTTTCTTGATGATGTG-3’ and reverse – 5’-ACCTCAACCAGAAAAACCAG-3’. PCR reaction conditions are as follows: 1.5 μL 10x iTaq PCR buffer, 1.2 μL MgCl_2_ (50 mM), 0.6 μL each primer (10 μM), 0.67 μL DMSO, 0.3 μL dNTPs (2.5 mM each), 0.25 μL iTaq polymerase (BioRad #170-8870), 8.38 μL water, and 2.5 μl gDNA per reaction. Thermocycler conditions were as follows: 94°C x 1’; 94°, 56°, 72° x 15”, 15”, 20” for two cycles; 94°, 54°, 72° x 15”, 15”, 20” for two cycles; 94°, 52°, 72° x 15”, 15”, 20” for two cycles; 94°, 50°, 72° x 15”, 15”, 20” for two cycles; 94°, 48°, 72° x 15”, 15”, 20” for two cycles; 94°, 46°, 72° x 15”, 15”, 20” for two cycles; 94°, 44°, 72° x 15”, 15”, 20” for two cycles; 94°, 42°, 72° x 15”, 15”, 20” for ten cycles; 94°, 40°, 72° x 15”, 15”, 20” for ten cycles; 72° x 2’; 4° infinite hold.

After PCR a restriction enzyme digest was performed on the PCR product to distinguish between the wild-type and F508del alleles. For each reaction, a mixture containing 2 μL NEBuffer1, 2.8 μL water, and 0.2 μL Rsa I (10 U/μl; New England BioLabs, #R0167S) was prepared and added to the tube containing the PCR product. The restriction digest reaction was incubated at 37 °C for one hour. Following digestion, the PCR product was run on at 3% agarose gel (1% agarose, GPG and 2% agarose, Supra Sieve/GPG; American Bioanalytical) in 1x Tris acetate/EDTA (TAE) buffer. The genotype was scored as follows: bands at 109 and 21 bp indicate a wild-type allele, while undigested bands at 130 bp indicate the mutant, F508del, allele.

### In utero injections

Time dated pregnant mice (8-12 weeks old) 15 or 16 days post conception were anesthetized with inhaled isoflurane (3% vol/vol for induction, 2% vol/vol for maintenance). The gravid uterus was exposed through a midline laparotomy incision. Lyophilized PNA/DNA nanoparticles were resuspended by vortex and water bath sonication in 1x dPBS to a concentration of 9 mg ml^-1^ (DiO NPs) or 12 mg ml^-1^ (PNA/DNA NPs). Intravascular injections were performed at E15.5 and intra-amniotic injections were performed at E16.5. A volume of 15 μl of NP suspension was drawn up into a glass micropipette (tip diameter ∼60 μm) and injected intravascularly via vitelline vein of each fetus using a pneumatic microinjector (Narishige; Japan). For intra-amniotic injections, 20 μl of NP suspension was injected directly into the amniotic cavity.

### Microscopy

Pregnant mice were euthanized three hours after in utero NP treatment. Fetuses were delivered and washed in 1x dPBS. Fetuses were fixed overnight in 4% paraformaldehyde (Electron Microscopy Sciences; Hartfield, PA) at 4 °C. The tissues were next dehydrated in 20% sucrose for 24 hours and embedded in Tissue-Tek Optimal Cutting Temperature (OCT) Compound (Torrance, CA). Frozen 15 μm fetal lung section were mounted on glass slides and stained with Hoescht dye. Confocal imaging of the frozen sections was performed on a Zeiss Axio Observer Z1 microscope (Oberkochen, Germany).

### Flow cytometry

For flow cytometry experiments, pregnant dams and fetuses were euthanized three hours after fluorescent NP or four days after PNA/DNA NP delivery. The lung was dissected from each fetus and washed in 1x dPBS. Single cell suspensions of lung tissue were made by homogenizing the organ through a 70 μm cell strainer (Falcon). Single cell suspensions of each organ were fixed in 4% paraformaldehyde. For the lung cellular tropism studies, single cell suspensions of lung cells were resuspended n 2% FBS in PBS and incubated with a 1:1000 dilution of FcR blocking reagent (Miltenyi Biotec Inc., Auburn, CA). The cells were subsequently stained for EpCAM, CD31, and CD45 using the following antibodies: EpCAM-Super Bright 436 (Cat# 62-5791-82, Invitrogen; Carlsbad, CA), CD31-APC (Cat# 17-0311-82, Invitrogen; Carlsbad, CA), and CD45R-PE-Vio770 (Cat# 130-102-817, Miltenyi Biotec Inc.; Auburn, CA). Nanoparticle delivery and uptake (fluorescent NPs) or eGFP expression (γPNA/DNA NPs in the 654-eGFP mouse) was measured using a BD FACSCalibur (BD Biosciences; Franklin Lakes, NJ). Flow cytometry data obtained was analyzed using FlowJo software (FlowJo, LLC; Ashland, OR). For the lung tropism studies the cell populations were defined as follows: endothelial – CD31+, EpCAM-, CD45-; epithelial CD31-,EpCAM+,CD45-, and immune CD31-, EpCAM-, CD45+^44,53^.

### Nanoparticle stability

Amniotic fluid was collected and pooled from C57BL/6 fetuses at E16.5. PLGA and PLGA/PBAE/MPG NPs were resuspended in dH_2_O or E16.5 murine amniotic fluid. DLS measurements of the hydrodynamic diameter and PDI were obtained in 10 minute intervals over an hour at 25°C using a Malvern Nano-ZS (Malvern Instruments, UK).

### Oligonucleotides

For experiments involving the eGFP mouse model, mini-PEG γPNA monomers were prepared from Boc-(2-(2-methoxyethoxy)ethyl)-L-serine as a starting material by a series of multistep synthetic procedures including reduction, mitsunobu reaction, nucleobase (A,C,G and T) conjugation and then ester cleavage^54^. At each step the respective product was purified by column chromatography^21^. PNA oligomers were synthesized on solid support using Boc chemistry^54^. The oligomers were synthesized on MBHA (4-methylbenzhydrylamine) resin according to standard procedures of Boc chemistry. A kaiser test was performed at each step to measure complete coupling and double coupling was performed if it was required. The oligomers were cleaved from the resin using an m-cresol/thioanisole/TFMSA/TFA (1:1:2:6) cocktail, and the resulting mixtures were precipitated with ethyl ether, purified by reversed phase-high-performance liquid chromatography (acetonitrile:water) and characterized with a matrix-assisted laser desorption/ionization time-of-flight mass spectrometer^18^. The sequence of γPNA used in this study is H-KKK-JTTTJTTTJTJT-OOO-T<u>C</u>T<u>C</u>T<u>T</u>T<u>C</u>T<u>T</u>T<u>C</u>A<u>G</u>G<u>G</u>C<u>A</u>-KKK-NH_2_. Underlined indicates γPNA residues; K, lysine; J, pseudoisocytosine; O, 8-amino-2,6,10-trioxaoctanoic acid linkers connecting the Hoogsteen and Watson-Crick domains of the tcPNA. The single-stranded donor DNA oligomer was prepared by standard DNA synthesis except for the inclusion of three phosphorothioate internucleoside linkages at each end to protect against nuclease degradation (Midland Certified Reagent Company; Midland, TX). The 60 bp donor DNA used in this study with the correcting nucleotide underlined is: 5’A(s)A(s)A(s)GAATAACAGTGATAATTTCTGGGTTAAGG<u>C</u>AATAGCAATATCTCTGCATATAAA(s)T(s)A(s)T3’.

The PNA used for CF studies binds in the murine CFTR gene and is as follows: H-KKK-JTTTTJJJ-OOO-CCCTTTTCAAGGTGAGTAG-KKK-NH_2_.

The murine CFTR donor DNA is as follows :5′T(s)C(s)T(s) TATATCTGTACTCATCATAGGAAACACCA<u>AAG</u>ATAATGTTCTCCTTGATAGTACC(s) C(s)G(s)G3′. The phenylalanine codon insertion base pair is underlined. As above, donor DNA was synthesized by Midland Certified Reagent Company (Midland, TX) and contains three phosphorothioate internucleoside linkages at both the 5’- and 3’-ends.

### Digital PCR

gDNA was extracted from tissues using the Wizard SV DNA Purification System (Promega, Madison, WI) according to manufacturer’s instructions. The concentration of extracted gDNA samples was measured using a QuBit® dsDNA BR assay kit (Invitrogen, Carlsbad, CA) according to manufacturer’s instructions. Up to 80 ng of gDNA was used for each sample per reaction. PCR reactions were set up as followed: 11 μl 2x ddPCR™ supermix for probes (no dUTP) (Bio-Rad, Hercules, CA), 0.2 μl forward primer (100 μM), 0.2 μl reverse primer (100 μM), 0.053 μl β-thal probe (100 μM), 0.053 μl wild-type probe (100 μM) (Integrated DNA Technologies, Coralville, IA), 0.5 μl EcoR1, 10 μl gDNA and dH_2_O. Droplets were generated using the Automated Droplet Generator (AutoDG™) (Bio-Rad). Thermocycling conditions for PCR for the eGFP mouse were as follows: 95°C 10 min, (94°C 30s, 55.3°C 1 min – ramp 2°C/s) x 40 cycles, 98°C 10 min, hold at 4°C. Thermycycling conditions for the CFTR PCR were as follows: 95°C 10 min, (94°C 30s, 53.7°C 1 min – ramp 2°C/s) x 40 cycles, 98°C 10 min, hold at 4°C. Droplets were allowed to rest at 4°C for at least 30 minutes after cycling and were then read using the QX200™ Droplet Reader (Bio-Rad). Data were analyzed using QuantaSoft™ software. Data are represented as the fractional abundance of the wild-type (control) or edited (β-globin or CFTR) allele. The primers used for ddPCR were as follows: eGFP intron forward: 5’-ACCATTCTAAAGAATAACAGTGA-3’, eGFP intron reverse: 5’-CCTCTTACATCAGTTACAATTT-3’, CFTR forward: 5’-TGCTCTCAATTTTCTTGGAT-3’, CFTR reverse: 5’-AAGCTTTGACAACACTCTTA-3’. The probes used for ddPCR were as follows: eGFP wild-type (FAM): 5’ TGGGTTAAGGCAATAGCAA, eGFP mutant (HEX): 5’ TCTGGGTTAAGGTAATAGCAAT, edit CFTR (FAM): 5’- CACCAAAGATAATGTTCTCCT-3’, F508del CFTR (HEX): 5’- ATCATAGGAAACACCAATGATAT-3’.

### Deep sequencing

Genomic DNA (gDNA) from the tissues of PNA/DNA NP treated CF mice was purified using the SV DNA Purification System (Promega, Madison, WI) according to manufacturer’s instructions. PCR reactions were performed with high fidelity TAQ polymerase (Invitrogen; Carlsbad, CA). Each PCR tube consisted of 28.2 μL dH_2_O, 5 µL 10x HiFi Buffer, 3 μL 50mM MgCl_2_, 1 μL dNTP, 1 μL each of forward and reverse primer, 0.8 μL High Fidelity Platinum Taq Polymerase and 10 μL 40 ng/ml gDNA. Thermocycler conditions were as follows: 94°C 2min (94°C 30s, 55°C 45s, 68°C 1 min) x35 cycles, 68°C 1 min, hold at 4°C. PCR products were purified using the QIAquick PCR Purification Kit (Qiagen; Hilden, Germany). PCR products were prepared by end-repair and adapter ligation according to Illumina protocols (San Diego, CA), and samples were sequenced by the Illumina HiSeq 2500 with 75 paired-end reads at the Yale Center for Genome Analysis. Samples were analyzed as previously described using the variant caller FreeBayes on basepairtech.com^55^. To be considered a variant, there had to be at least two observations of that change in the amplicon, and the change had to be present at a frequency of at least 0.002 (set as the minimum alternative fraction) (0.2%). Reads for samples at any one base were capped at 10,000 randomly selected reads. The analysis pipeline was developed by basepairtech.com. The primers used for murine CFTR were as follows: forward primer: 5’-TCTGCTCTCAATTTTCTTGGA-3’; reverse primer: 5’- GGCAAGCTTTGACAACACTC-3’. Primers for off-target effects (forward listed first) : OT- 00 – 5’-TTGTCAAAGCTTGCCAACTA-3’, 5’- ACGGTATCATCCCTGAAAAG-3’; OT-01 – 5’- TACCTTCGGTATCCCAAATCTC-3’, 5’- GCCTGTGATATGATAGACACCT-3’; OT-02–5’- GAGCCTACTGGGAGGTAAAAT-3’, 5’- GGACCTGATTACCTTGGGTAT-3’, OT-03–5’- ATGTGAGAGGACTCTGTGAA-3’, 5’- ACCTGTACTGGTTTATAGGG-3’; OT-04 – 5’- TATCACATTGGCCATCTCAG-3’, 5’- GGTACAAGGATAGCAGTAGC-3’; OT-05 – 5’- TGGTACAAGGATGGCAGTA-3’, 5’- CCATTACCTCGGGAAGATTT-3’; OT-06 – 5’- AATGCCCAATACAACAGATTT-3’, 5’- GAGCCATCTTTTGATGTTCAG-3’; OT-07 – 5’- CCTGACTGATGGATGACGAGTTA-3’, 5’- TCAGTCCTGGTTGGAAAAGC-3’; OT-08 – 5’- CCTCACCTTAACGAGCAAA-3’, 5’- TCAATGGACTCTCCCTAGAC-3’; OT-09 – 5’- TGGTACAAGGATGGCAGTA-3’, 5’- GGATCTTCTTGGCTATCACA-3’; OT-10 – 5’-GTCTCAGTCCTGGTAGGAAA-3’, 5’- AATACCCTACTGCCCTACTC-3’; OT-11 – 5’- TGGATCTTCCTGGTGATTTTG-3’, 5’- TTATAAATTTCCCAGACTAGGCTATAA-3’^11^.

### RNA-seq

Total RNA was prepared from the lungs of three adult mice (8 weeks old) and three E16.5 fetal mouse lungs using the Qiagen RNeasy kit (Hilden, Germany) per manufacturer’s instructions. PolyA selection, library preparation, and sequencing (75 bp paired-end reads) was performed by the Yale Center for Genome Analysis using the Illumina HiSeq 2500. Analysis was performed by Francesc Lopez-Giraldez (Yale Center for Genome Analysis).

### Functional analysis – nasal and rectal potential difference, Ussing chamber, BAL fluid

Nasal and rectal potential differences (NPD, RPD) were measured as previously described^56^. Briefly, mice were anesthetized with ketamine/xylazine, and one electrode probe placed into one nostril (NPD) or rectum (RPD), with a reference electrode with 3% agar in Ringer’s solution placed subcutaneously. A microperfusion pump was used to flow solution through the electrode probe at 0.2 mL hour^-1^ for NPD and 0.5 mL hour^-1^ for RPD. Potential differences were measured first with a control Ringer’s solution, then with Ringer’s solution containing 100 μM amiloride, then a chloride-free solution with amiloride, and then with forskolin and IBMX. NPDs were measured once the mice reached a weight of at least 20g.

For ex vivo BAL fluid and Ussing chamber analyses, mice were euthanized eight months after in utero NP treatment and animals were heart-perfused with heparinized dPBS. To collect BAL fluid samples lungs were filled with 2 mL of 1x dPBS containing protease inhibitors qnd 0.5M EDTA at pH 8^11^. This fluid was collected for cell count. For Ussing analyses, samples of epithelia from the duodenum, jejunum, ileum, distal colon, and rectum were mounted in Ussing chambers and analyses were performed as described by Grubb^57^. The 0 Cl-solution used for all GI tract tissues contained 5 mM barium hydroxide to block potassium currents^11^.

### Statistical analysis

Results were analyzed using GraphPad Prism. Unless otherwise stated, data are presented as individual data points ± s.e.m. Data are compared by one-way or two-way ANOVA with repeated measures when appropriate. Turkey’s posttest was used to correct for multiple comparisons

## Supplementary Figures

**Supplementary Table 1.**
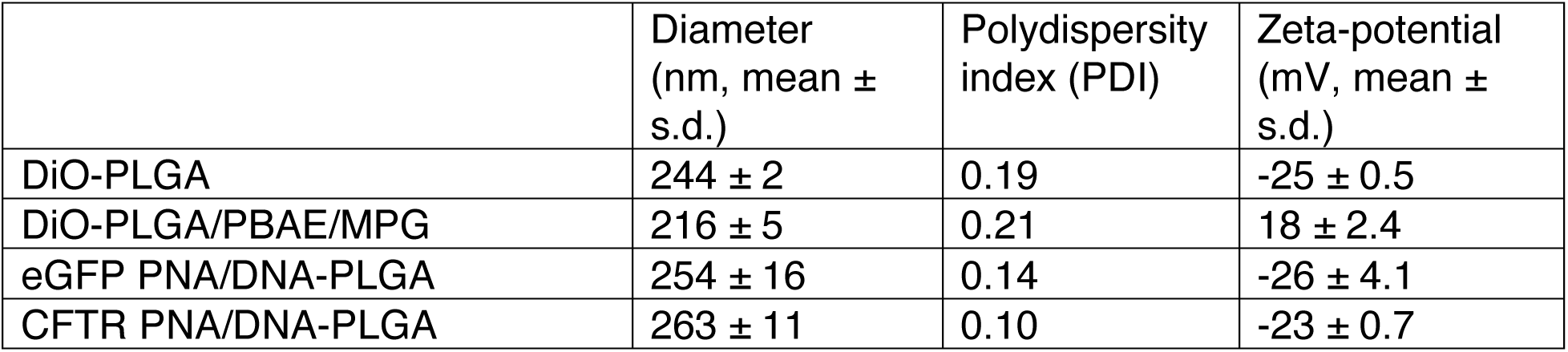
Nanoparticle characterization.

| | Diameter<br>(nm, mean ±<br>s.d.) | Polydispersity<br>index (PDI) | Zeta-potential<br>(mV, mean ±<br>s.d.) |
| --- | --- | --- | --- |
| DiO-PLGA | 244 ± 2 | 0.19 | -25 ± 0.5 |
| DiO-PLGA/PBAE/MPG | 216 ± 5 | 0.21 | 18 ± 2.4 |
| eGFP PNA/DNA-PLGA | 254 ± 16 | 0.14 | -26 ± 4.1 |
| CFTR PNA/DNA-PLGA | 263 ± 11 | 0.10 | -23 ± 0.7 |

**Supplementary Figure 1.**
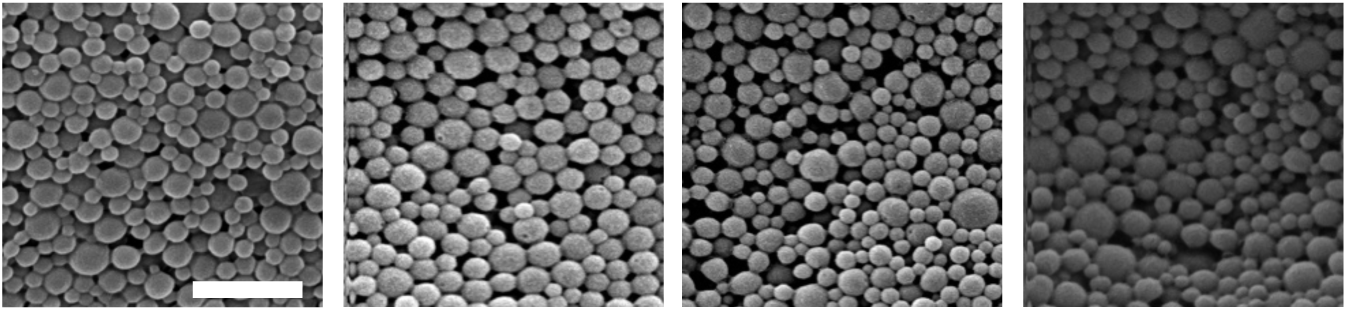
Scanning electron micrographs of DiO-PLGA, DiO-PLGA/PBAE/MPG, eGFP PNA/DNA-PLGA, and CFTR PNA/DNA-PLGA (left to right) nanoparticles, scale bar = 1 μm.

**Supplementary Figure 2.**
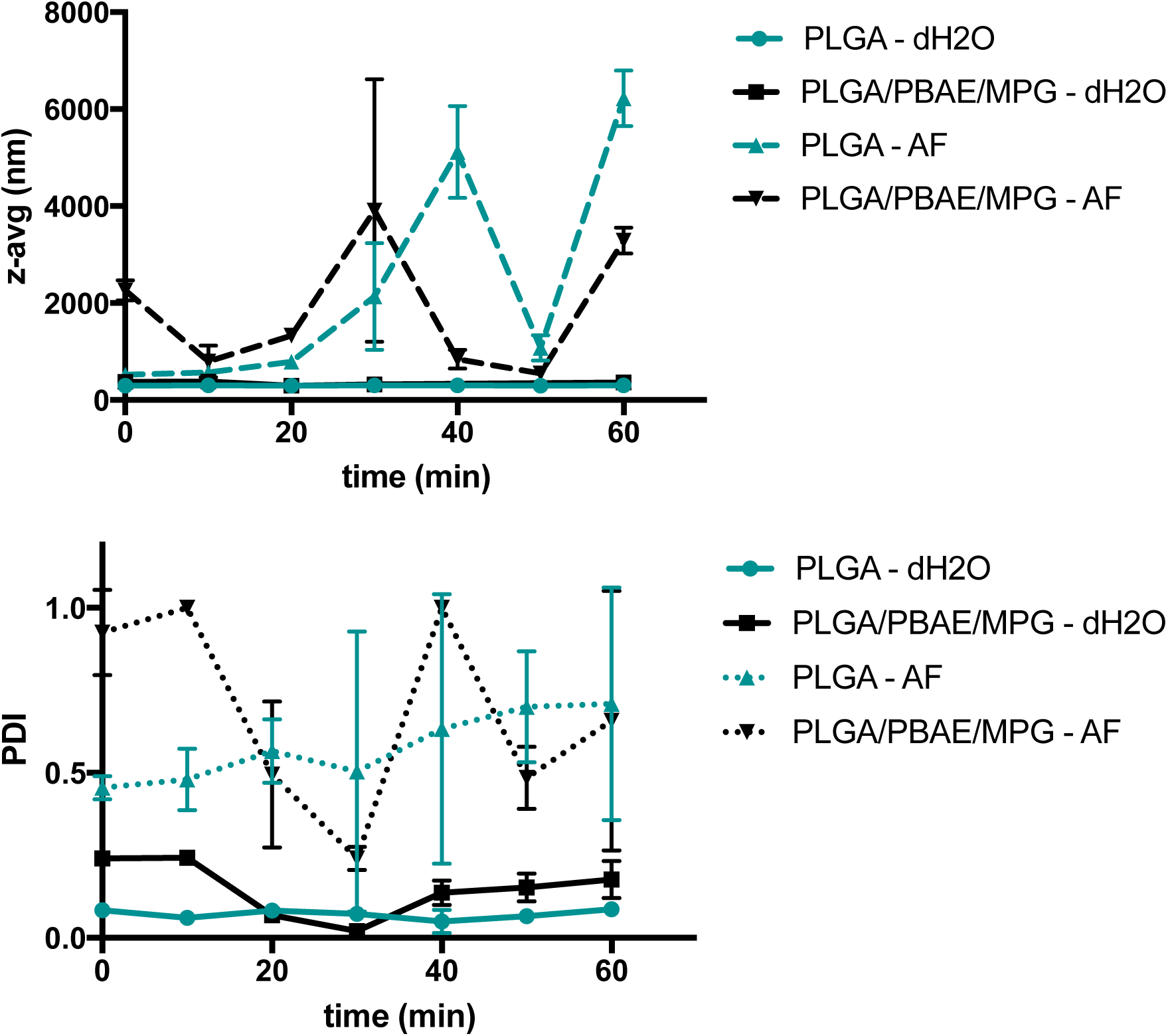
Nanoparticle (NP) stability in murine amniotic fluid. The hydrodynamic diameter (z-avg, top) and polydispersity index (PDI, bottom) were measured for both PLGA (teal) and PLGA/PBAE/MPG NPs (black) resuspended in dH_2_O (solid lines) or E16.5 murine amniotic fluid (AF, dashed lines) every ten minutes for an hour.

**Supplementary Figure 3.**
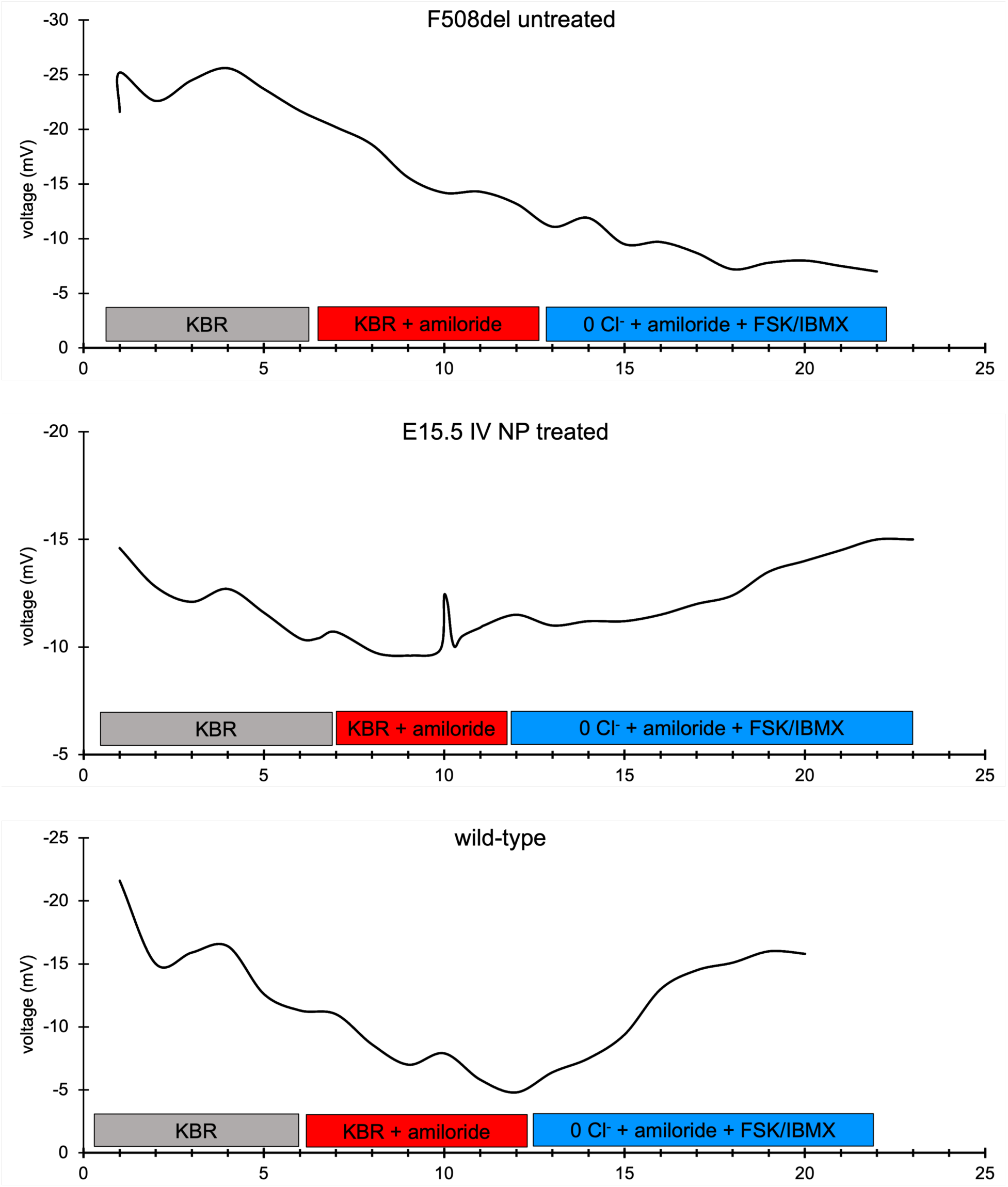
Representative nasal potential difference of a CF mouse four months after IV PNA nanoparticle treatment compared to an untreated F508del mouse and a wild-type mouse. The F508del CF mouse nasal epithelia have a large lumen-negative potential that is amiloride-sensitive and lack forskolin stimulated chloride efflux (top). Amiloride is an inhibitor of the epithelial sodium channel (ENaC). CFTR downregulates ENaC, which is why affected CF tissue is more sensitive to amiloride. Forskolin increases levels of cAMP by stimulating adenylate cyclase; and IBMX inhibits phosphodiesterase, which also increases intracellular cAMP; increased cAMP stimulates chloride transport. The IV treated mice (middle) display a response similar to wild-type in which the epithelia display a more modest amiloride-sensitive response (ENaC downregulation) and a strong cAMP-stimulated chloride efflux (surrogate for CFTR activity).

**Supplementary Figure 4.**
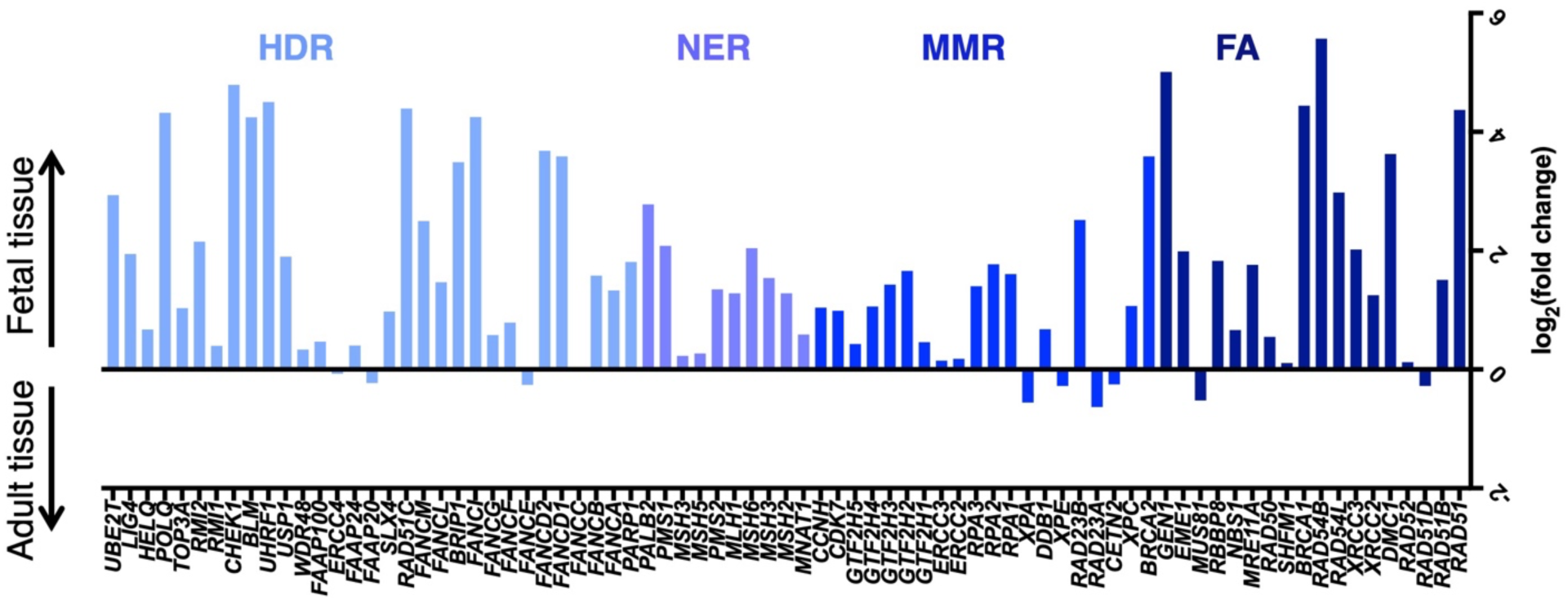
DNA repair pathways are elevated in the fetal lung. RNAseq analysis of genes in DNA repair pathways involved in gene editing. Each color represents clusters of genes associated with different repair pathways, from the left (light blue) to the right (navy blue): homology directed repair (HDR), nucleotide excision repair (NER), mismatch repair (MMR), and Fanconi anemia (FA).

**Supplementary Figure 5.**
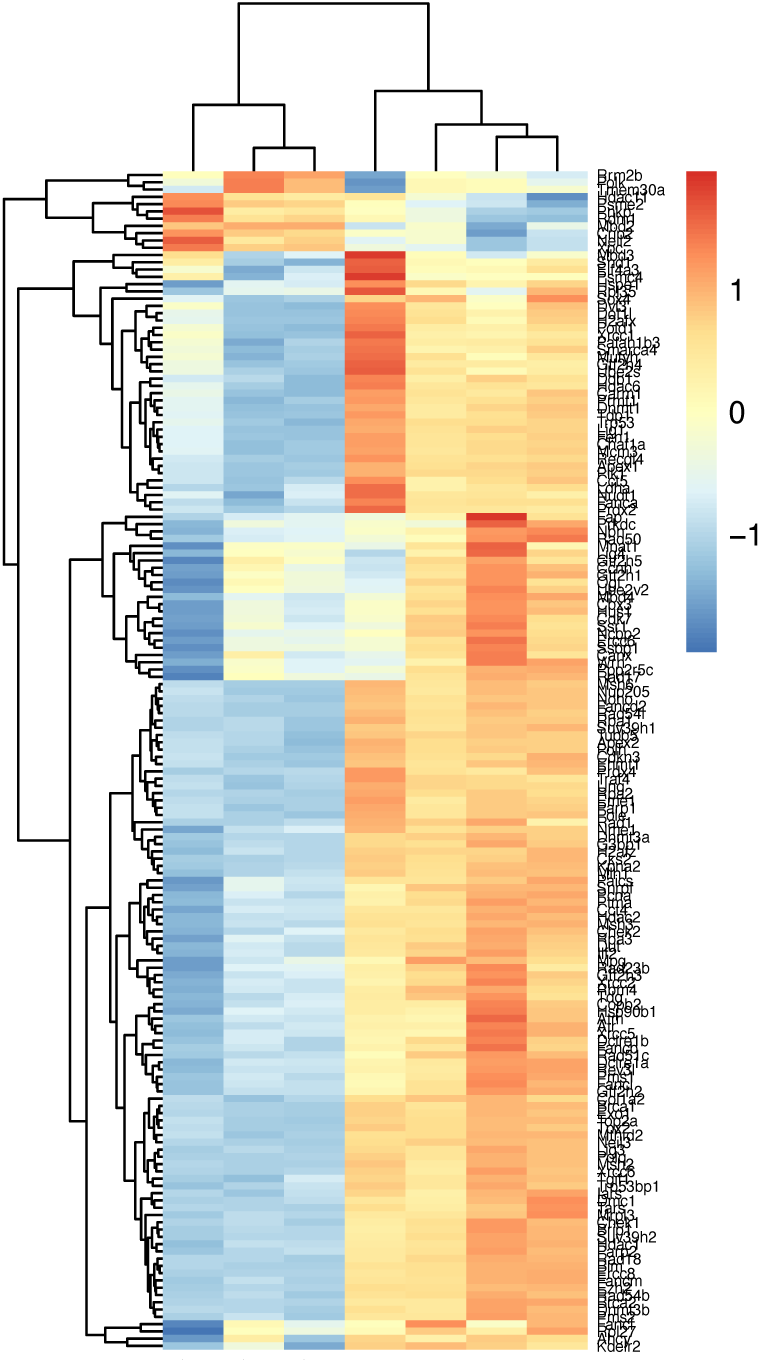
Heat map of DNA repair gene expression in adult and fetal lung. Of 226 DNA repair genes analyzed, 162 displayed significant differences in expression between the fetal and adult lungs, with the majority of DNA repair genes having higher expression in fetal lung tissue. Adult lung samples are grouped on the left and fetal lung samples are grouped on the right.

**Supplementary Table 2.**
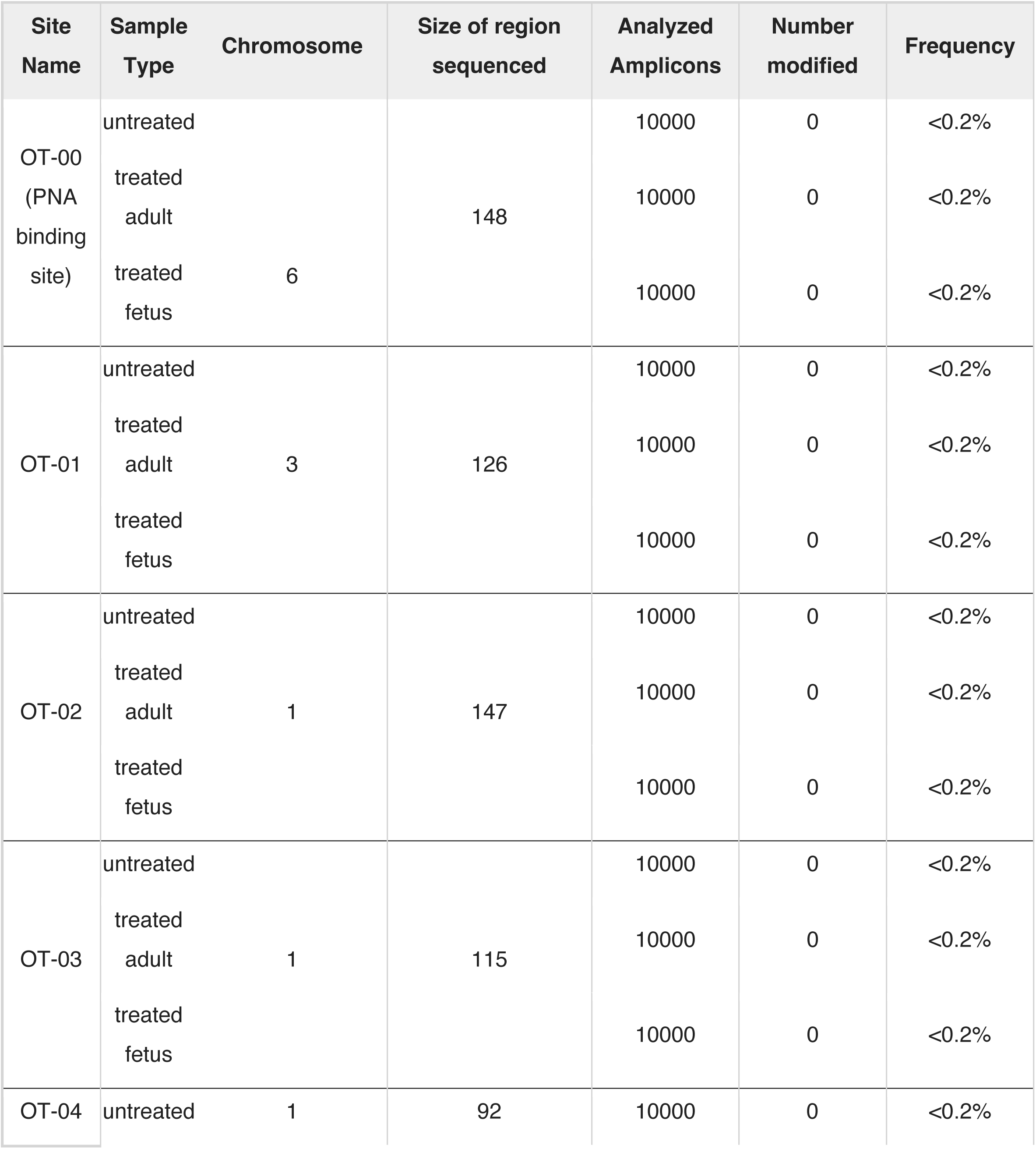
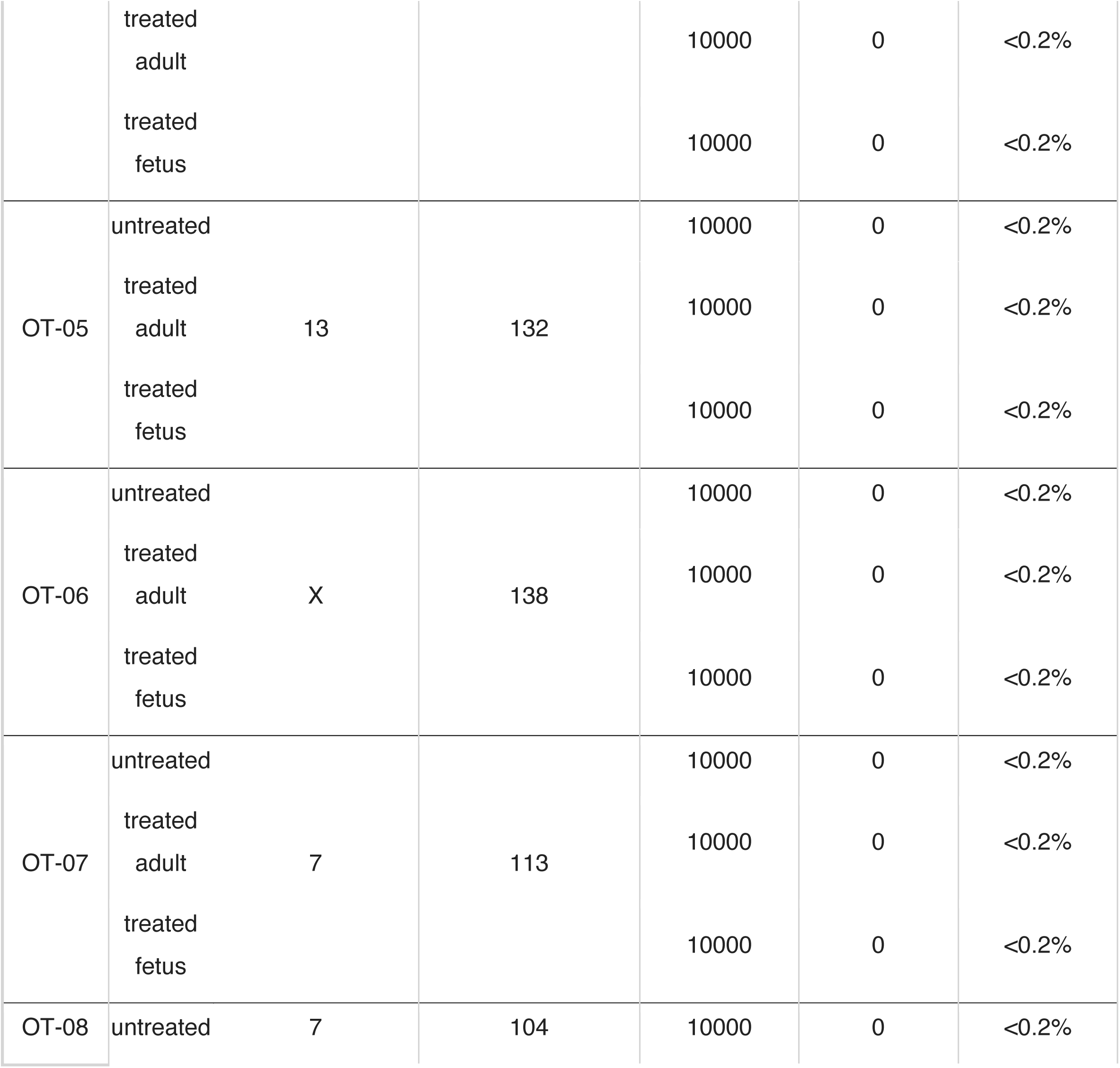
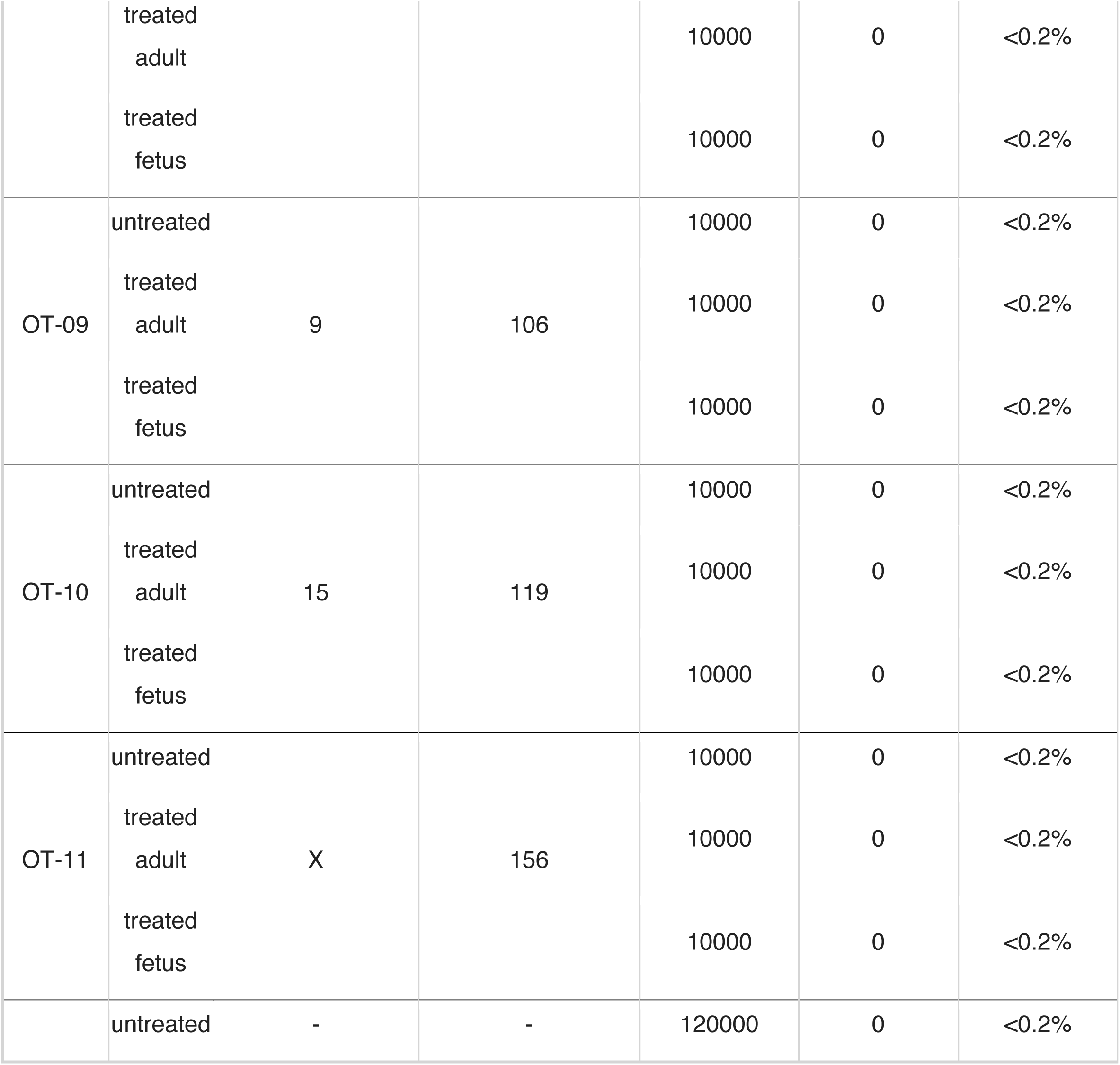
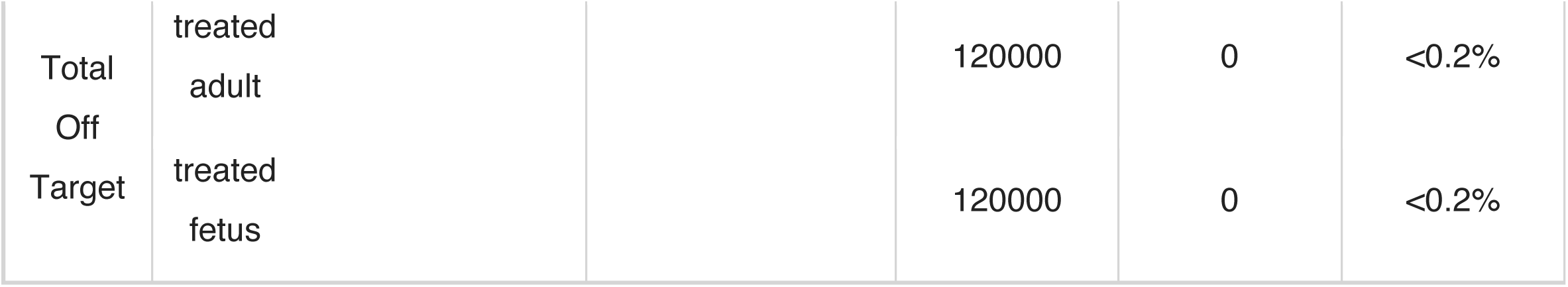
Deep sequencing off-target (OT) analysis of lung genomic DNA from CF mice that were untreated, fetal mice four days after IV in utero PNA NP treatment, or adult mice eight months after IV in utero PNA NP treatment at the PNA binding site (OT-00) and eleven genomic sites with partial PNA binding site (OT-01 – OT-10) or donor DNA (OT-11) homology. The size of the region sequenced around each site, the total number of amplicons sequenced and the number of amplicons with modified sequences are listed.

## Acknowledgements

We thank Emanuela Bruscia for invaluable conversations and Trucode Gene Repair Inc. for providing a portion of the PNAs used in this study. This work was supported by NIH grant R01 HL125892 (to P.M.G., W.M.S., and M.E.E.), NIH grant UG3 HL147352 (to P.M.G. and W.M.S.), NIH grant R01 EB032791 (to W.M.S. and D.H.S.), NIH grant F30 HL134252 (to A.S.R.), NIH grant K99/R00 HL151806 (to A.S.P.-D.), NIGMS Medical Scientist Training Program T32GM07205 (to A.S.R. and E.Q.), and Cystic Fibrosis Foundation grants EGANXX15, EGANXX19 (to P.M.G., W.M.S., and M.E.E.), PIOTRO20F0, PIOTRO21F5 (to A.S.P.-D.), and STITEL17G0 (to P.M.G., W.M.S., M.E.E. and D.H.S.).

## Author Contributions

A.S.R., R.P., A.S.P.-D., D.H.S., P.M.G., W.M.S., and M.E.E. designed the experiments. C.B., R.P., E.Q., A.G., R.N., H.M., M.R.F.-W., V.L., S.A., J.F., and D.H.S. performed the experiments. A.S.R., R.P., F.LG., and M.E.E. analyzed the data and prepared the figures. A.S.R. wrote the original draft of the manuscript. A.S.R., D.H.S., P.M.G., W.M.S., and M.E.E. edited the manuscript.

## Competing Interests

During the time this work was performed, A.S.R., E.Q., A.S.P.-D., P.M.G., W.M.S., and M.E.E. were consultants to Trucode Gene Repair Inc. A.S.R., E.Q., A.S.P.-D., P.M.G., W.M.S., D.H.S. and M.E.E. are inventors on patents and patent application related to this work. M.E.E., W.M.S., and A.S.P.-D. are cofounders of Xanadu Bio. W.M.S. is a member of the Board of Directors of Xanadu Bio and a consultant to Xanadu Bio, B3 Therapeutics, Stradefy Biosciences, Johnson & Johnson, Celanese, Cranius, and CMC Pharma.

